# Updates to the Alliance of Genome Resources Central Infrastructure Alliance of Genome Resources Consortium

**DOI:** 10.1101/2023.11.20.567935

**Authors:** Suzanne A. Aleksander, Anna V. Anagnostopoulos, Giulia Antonazzo, Valerio Arnaboldi, Helen Attrill, Andrés Becerra, Susan M. Bello, Olin Blodgett, Yvonne M. Bradford, Carol J. Bult, Scott Cain, Brian R. Calvi, Seth Carbon, Juancarlos Chan, Wen J. Chen, J. Michael Cherry, Jaehyoung Cho, Madeline A. Crosby, Jeffrey L. De Pons, Peter D’Eustachio, Stavros Diamantakis, Mary E. Dolan, Gilberto dos Santos, Sarah Dyer, Dustin Ebert, Stacia R. Engel, David Fashena, Malcolm Fisher, Saoirse Foley, Adam C. Gibson, Varun R. Gollapally, L. Sian Gramates, Christian A. Grove, Paul Hale, Todd Harris, G. Thomas Hayman, Yanhui Hu, Christina James-Zorn, Kamran Karimi, Kalpana Karra, Ranjana Kishore, Anne E. Kwitek, Stanley J. F. Laulederkind, Raymond Lee, Ian Longden, Manuel Luypaert, Nicholas Markarian, Steven J. Marygold, Beverley Matthews, Monica S. McAndrews, Gillian Millburn, Stuart Miyasato, Howie Motenko, Sierra Moxon, Hans-Michael Muller, Christopher J. Mungall, Anushya Muruganujan, Tremayne Mushayahama, Robert S. Nash, Paulo Nuin, Holly Paddock, Troy Pells, Norbert Perrimon, Christian Pich, Mark Quinton-Tulloch, Daniela Raciti, Sridhar Ramachandran, Joel E. Richardson, Susan Russo Gelbart, Leyla Ruzicka, Gary Schindelman, David R. Shaw, Gavin Sherlock, Ajay Shrivatsav, Amy Singer, Constance M. Smith, Cynthia L. Smith, Jennifer R. Smith, Lincoln Stein, Paul W. Sternberg, Christopher J. Tabone, Paul D. Thomas, Ketaki Thorat, Jyothi Thota, Monika Tomczuk, Vitor Trovisco, Marek A. Tutaj, Jose-Maria Urbano, Kimberly Van Auken, Ceri E. Van Slyke, Peter D. Vize, Qinghua Wang, Shuai Weng, Monte Westerfield, Laurens G. Wilming, Edith D. Wong, Adam Wright, Karen Yook, Pinglei Zhou, Aaron Zorn, Mark Zytkovicz

## Abstract

The Alliance of Genome Resources (Alliance) is an extensible coalition of knowledgebases focused on the genetics and genomics of intensively-studied model organisms. The Alliance is organized as individual knowledge centers with strong connections to their research communities and a centralized software infrastructure, discussed here. Model organisms currently represented in the Alliance are budding yeast, *C. elegans*, *Drosophila*, zebrafish, frog, laboratory mouse, laboratory rat, and the Gene Ontology Consortium. The project is in a rapid development phase to harmonize knowledge, store it, analyze it, and present it to the community through a web portal, direct downloads, and APIs. Here we focus on developments over the last two years. Specifically, we added and enhanced tools for browsing the genome (JBrowse), downloading sequences, mining complex data (AllianceMine), visualizing pathways, full-text searching of the literature (Textpresso), and sequence similarity searching (SequenceServer). We enhanced existing interactive data tables and added an interactive table of paralogs to complement our representation of orthology. To support individual model organism communities, we implemented species-specific “landing pages” and will add disease-specific portals soon; in addition, we support a common community forum implemented in Discourse. We describe our progress towards a central persistent database to support curation, the data modeling that underpins harmonization, and progress towards a state-of-the art literature curation system with integrated Artificial Intelligence and Machine Learning (AI/ML).

## Introduction

As has been discussed at length elsewhere (e.g., Oliver et al. 2016; Wood et al., 2022), model organism knowledgebases (aka model organism databases; MODs) provide daily utility to researchers for the design and interpretation of experiments, to computational biologists for curated datasets, and to genomic researchers for annotated genomes. Some of the major uses of the MODs have been one-stop shopping for all information about a particular gene or obtaining cleansed datasets with standard metadata for computational analyses.

The Alliance of Genome Resources (referred to herein as the Alliance) is a consortium of MODs and the Gene Ontology Consortium (GOC). The mission of the Alliance is to support comparative genomics as a means to investigate the genetic and genomic basis of human biology, health, and disease. To promote sustainability of the core community data resources that make up the Alliance, we implemented an extensible “knowledge commons” platform for comparative genomics built with modular, re-usable infrastructure components that can support informatics resource needs across a wide range of species (Alliance of Genome Resources, 2022; Howe et al., 2018; Bult and Sternberg, 2023). In 2022, the Alliance was recognized as a Core Global Biodata Resource by the Global Biodata Coalition (Anderson et al 2017).

Specifically, the Alliance of Genome Resources is organized as two interdependent units: Alliance Central and the Alliance Knowledge Centers. ***Alliance Central*** is responsible for developing and maintaining the software for data access and for the coordination of data harmonization and data modeling activities across our members. A primary goal of Alliance Central is to reduce redundancy in systems administration and software development for model organism knowledgebases and to deploy a unified ‘look and feel’ for access to, and display of, common data types and annotations across diverse model organisms and human, following Findability, Accessibility, Interoperability, and Reuse (FAIR) guiding principles. Model organism-specific knowledgebases serve as ***Alliance Knowledge Centers.*** Knowledge Centers are responsible for expert curation and submission of data to Alliance Central using Alliance Central infrastructure. Knowledge Centers also are responsible for organism-specific user support activities and for providing access to data types not yet supported by Alliance Central. The founding Alliance Knowledge Centers are *Saccharomyces* Genome Database (Engel et al. 2022), WormBase (Davis et al. 2022), FlyBase (Gramates et al 2022), Mouse Genome Database (Ringwald et al. 2022), the Zebrafish Information Network (Bradford et al. 2023), Rat Genome Database (Vedi et al 2023), and the Gene Ontology Consortium (Gene Ontology Consortium 2023). The newest member, Xenbase (Fisher et al, 2023), joined the Alliance consortium in 2022.

Here we describe our progress toward harmonizing information provided by our member resources, our development of a software infrastructure for ingest, curation, storage, analysis, and output of such information, and development of an efficient literature curation system. We also describe new features in our web portal at AllianceGenome.org.

## Community Homepages

The Alliance website features landing pages for each model organism in the Alliance consortium. These pages are accessed from the “Members” drop-down menu in the header on every Alliance page. These pages feature MOD-specific-content such as meetings, news, and other MOD-specific resource links. A common template allows users to find the same types of information in each landing page (**Figure 1**). As MODs transition their data and web services to the Alliance, their member pages will evolve into portals hosting additional MOD-specific data, tools, and links to organism-specific resources and will also accommodate the many unique data and tools from individual MODs.

**Figure 1.**
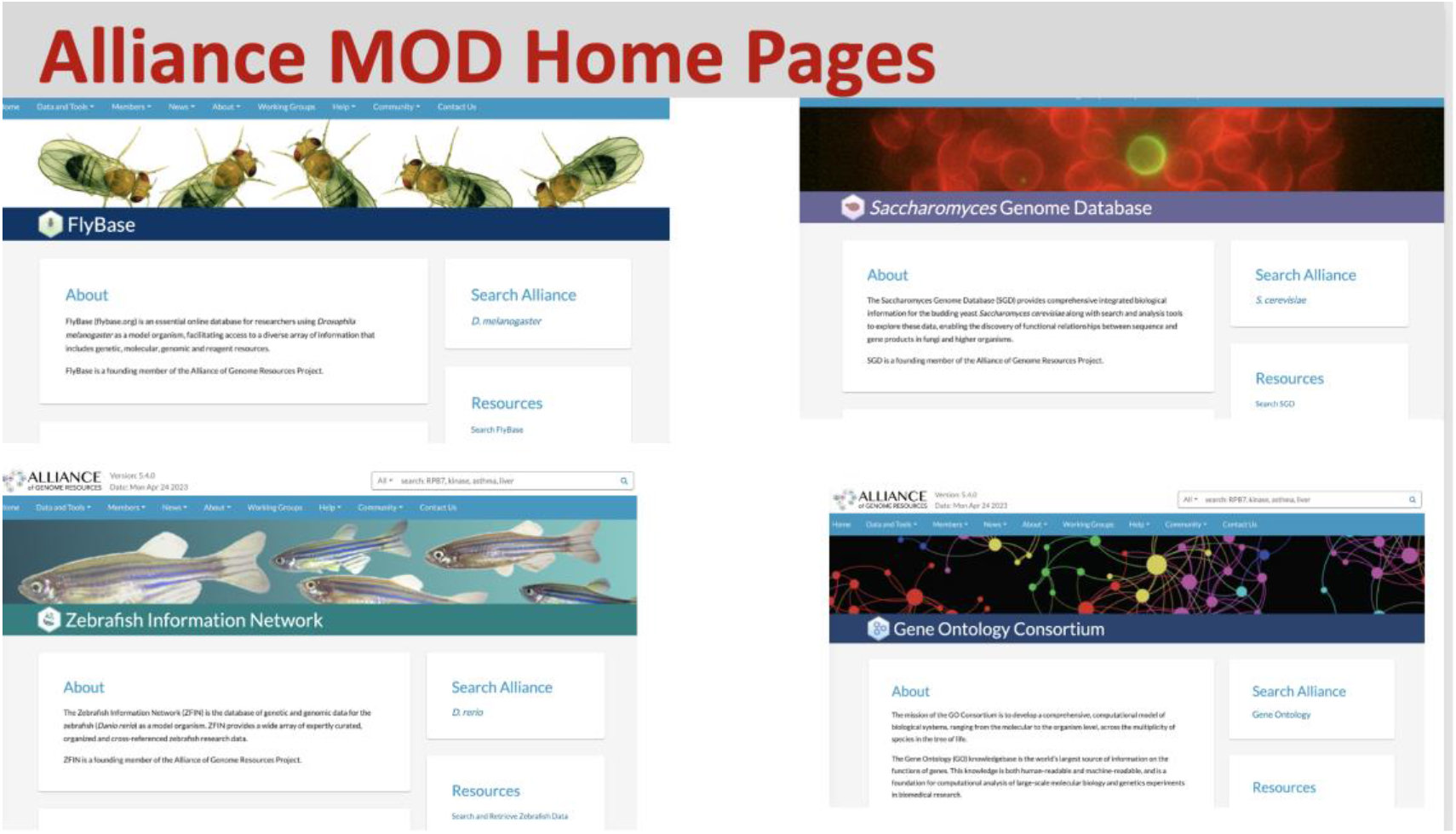
MOD landing pages at the Alliance Portal. A common look and feel that allows community-specific content.

## Paralogy

Gene pages include a new Paralogy section populated with data from the Drosophila Research & Screening Center (DRSC) Integrative Ortholog Prediction Tool (DIOPT) version 9.1 developed by the DRSC (Hu et al 2011, 2020). The assembly of protein sets and algorithmic inferences of their orthology from various sources was first centralized by the DRSC and then exported to the Alliance Central. We include the same data sources used for orthology, when these resources also provide paralogy information. Specifically, these resources have performed well on the standardized benchmarking from the Quest for Orthologs (QfO) Consortium (Nevers et al. 2022). Orthologous Matrix (OMA) (Altenhoff et al 2021) and PANTHER (Thomas et al. 2022) datasets were retrieved through the QfO benchmark portal (https://orthology.benchmarkservice.org), and Compara data were acquired directly from the EBI Compara FTP site. In addition, the DRSC conducted local analyses using Inparanoid (Persson and Sonnhammer. 2022), OrthoFinder (Emms and Kelly 2019), OrthoInspector (Nevers et al. 2019), and sonicParanoid (Cosentino and Iwasaki 2019) using a UniProt 2020 reference proteome. Direct data submissions from PhylomeDB (Fuentes et al. 2022) and the *Saccharomyces* Genome Database (SGD; Engel et al. 2022) were also integrated into the dataset.

The paralogy section is composed of a table (**Figure. 2**), similar to the orthology table, which contains the gene symbol of related paralogs, a calculated rank, alignment length as the number of aligned amino acids, percentage of similarity and identity, and a count of the algorithms or methods which call the paralogous match. The ranking score was developed to sort the paralogs by overall similarity, and was reviewed by curators to display optimally an acceptable rank order for well-studied sets of paralogs. The ranking score considers several factors, including alignment length, percent identity, and the number of paralogy methods that identify the paralog. Additional Information for rank determination and alignment length are available to the users via a clickable help icon located next to those column headers.

**Figure 2.**
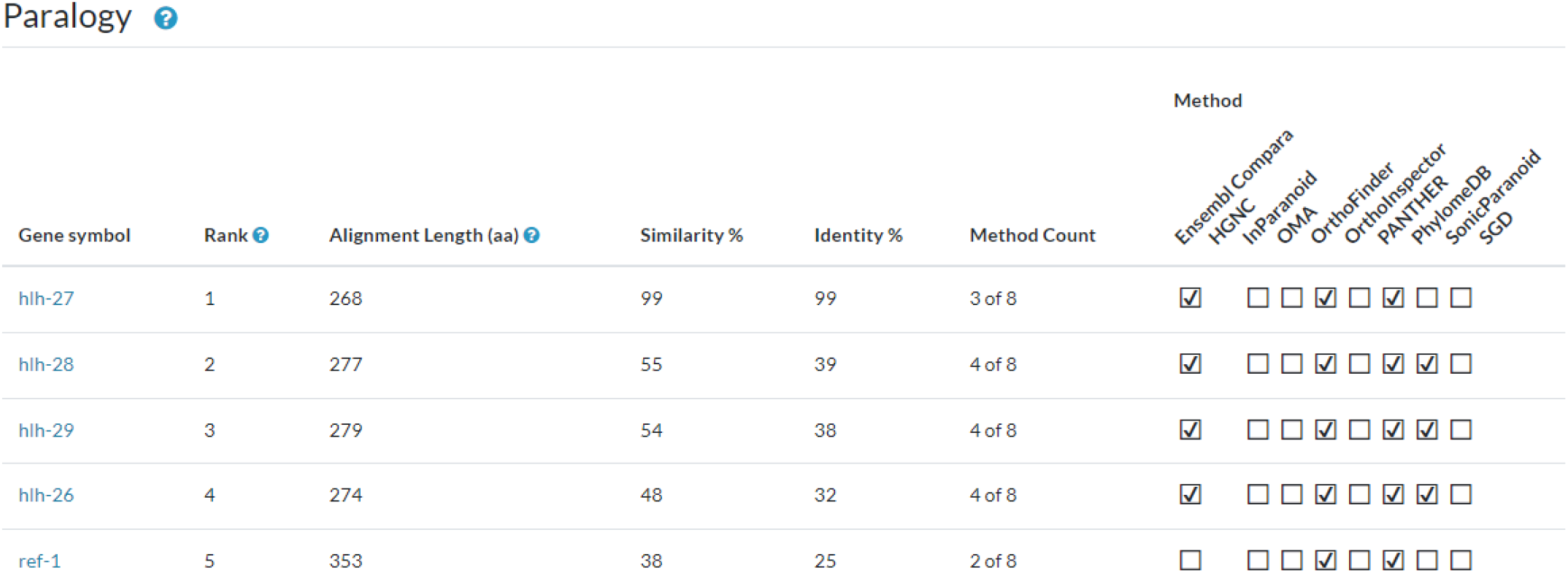
Paralog table for *C. elegans hlh-25*. The table presents a ranking of paralogs for the *hlh-25* gene, based on a weighted scoring algorithm that incorporates sequence conservation metrics. It lists the gene symbols, provides the alignment length in amino acids, and quantifies the similarity and identity percentages of genes paralogous to *hlh-25*. The methodology count, indicating the number of algorithms supporting the paralogous relationship, is also included. In this ranking, *hlh-27* is identified as the primary paralog due to its high similarity and identity scores, despite being recognized by fewer methods than *hlh-28*.

The paralog section was released with Alliance version 6.0.0. Future updates will include the ability to sort and filter the table by column values and the availability of these data via our bulk downloads page. The existing tables on the gene pages for Function, Disease, and Expression all contain checkboxes for “Compare Ortholog Genes” that allow users to search across species for these features. We will add the additional checkbox, “Compare Paralog Genes” to provide similar functionality for paralogous genes in a future Alliance release.

## Xenopus in the Alliance

Xenbase (Fisher et al 2023), the *Xenopus* knowledgebase, is the first knowledgebase to join the Alliance since the founding members initiated the consortium. *Xenopus* is an amphibian frog species used extensively in biomedical research, and in particular for experimental embryology, cell biology, and disease modeling with genome editing (Carotenuto et al., 2023; Kostiuk and Khokha, 2021). As a non-mammalian air-breathing tetrapod, *Xenopus* represents a valuable evolutionary transition between rodents and zebrafish for comparative genomic studies.

Xenbase is a large-scale knowledgebase built on a Chado schema foundation, so has design features related to Alliance foundational members. As a model system, two different *Xenopus* species are used interchangeably; *X. tropicalis* is a diploid that is the preferred system for genome editing and genetics, whereas *X. laevis* is an allotetraploid preferred for use in cell biology studies, microinjection, and microsurgery-style experimentation. *X. tropicalis* has 1:1 relationships between most genes and human orthologs (excluding paralogs) (Mitros et al., 2019),whereas *X. laevis* has two copies of most human orthologs. The allotetraploid formed via hybridization of two different frog species (Session et al 2016), and the complexities of genome evolution that subsequently occurred increase the difficulty of identifying orthology of the two *X. laevis* genes to their diploid relatives, including humans. Mapping of the diploid *X. tropicalis* genes to their human orthologs was performed with DIOPT, similarly to other model organisms in the Alliance. Because this method does not yet work in the context of an allotetraploid, the Alliance imports the *X. tropicalis* to *X. laevis* paralogy mappings from Xenbase, where they have been established through a combination of synteny analysis and manual curation. Dealing with how to incorporate the two new species with this ploidy complexity was one of the major challenges of adding *Xenopus* to the Alliance.

On the Xenbase side, as the new member team, Xenbase staff created exporters to upload content, on a regular schedule, formatted in a manner defined by the Alliance data ingest schema and using the Alliance File Management System and API access keys. Currently these data include orthology, the *Xenopus* anatomical ontology, standard gene information, gene expression data, publications, GO term associations, disease associations, anatomical phenotypes, genome details in the Alliance browser, and BLAST capacity. *Xenopus* genes can be found using the Alliance landing page search tool with *Xenopus* genes flagged by Xtr and Xla notations. The two copies of the genes in *X. laevis*, the allotetraploid, are further tagged as ‘*(symbol).L*’ and ‘*(symbol).S*’ to denote the genes on the long (L) and short (S) chromosome pairs of this species (e.g., *pax6.L* and *pax6.S*).Alliance release 6.0.0 has Xenbase data for 54,000 genes, 19,000 disease associations, over 45,000 gene expression records and more than 7,000 anatomical phenotypes. Expression and phenotype data will be available soon.

In addition to the rich data made available to the Alliance from *Xenopus* research, this effort also served as a valuable test case for understanding the level of effort and complexities engendered in the addition of new knowledgebases to the Alliance, and the functionality and adaptability of ingest system components.

## JBrowse sequence detail widget

Delivered in the recent Alliance 6.0.0 release, the “Sequence Detail’’’ section of all gene pages now uses JBrowse and javascript libraries to display an interactive widget that allows users to download DNA and amino acid sequences of genes in several possible configurations: genomic sequence highlighted with UTR, coding and intronic regions, CDS regions, and translated protein for example (**Figure 3**). We will extend the functionality of the widget variant detail pages, where both the wild-type and variant sequences will be provided. When the variant occurs in the context of a protein coding gene, changes to the coding sequence and resulting translated protein will also be displayed and available for download.

**Figure 3.**
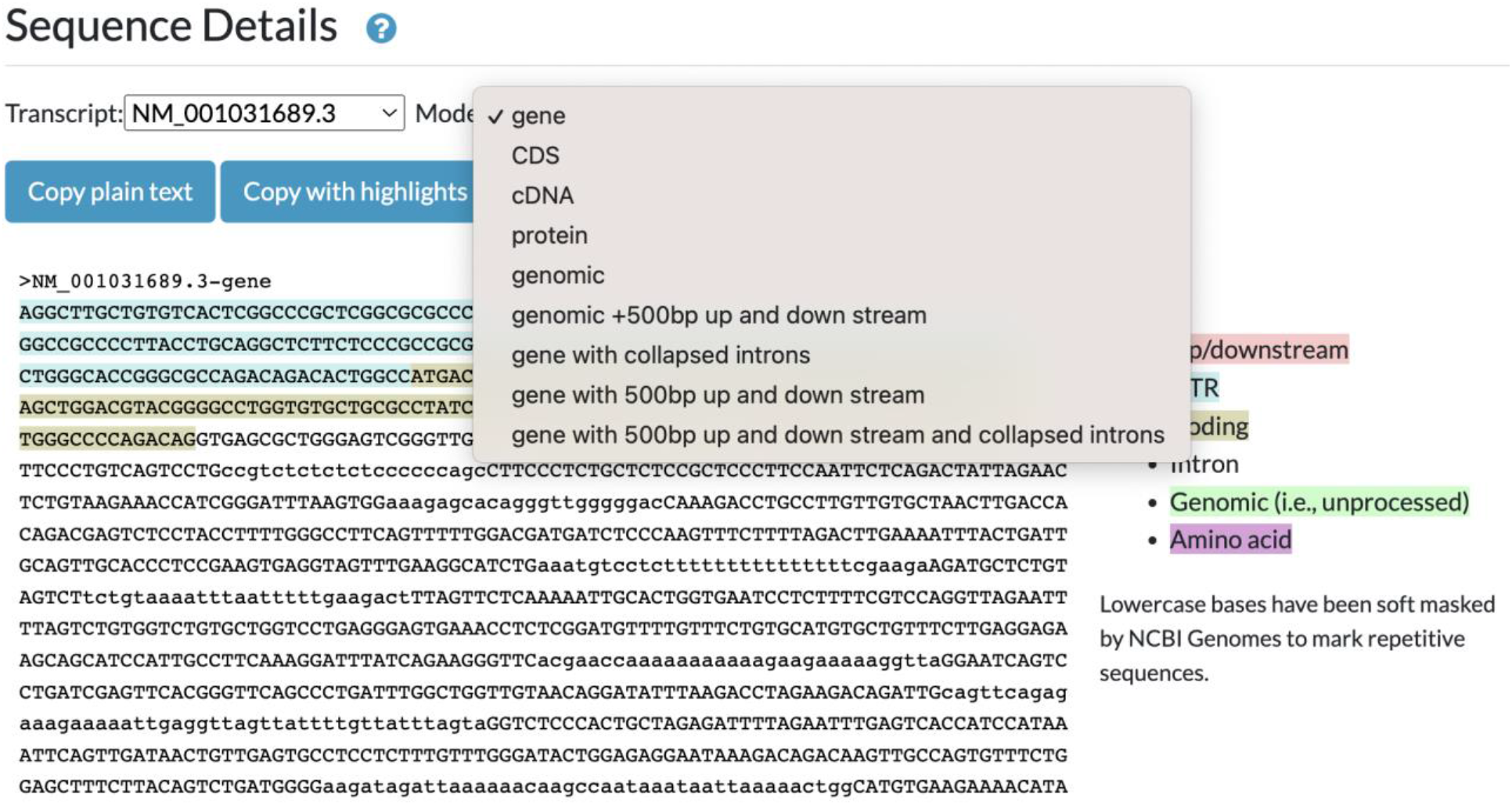
Sequence detail widget. Chosen views of a specific gene are readily available for copying as plain text or with highlights. 5’ region of the human PLAA gene.

## Model organism BLAST

For more than two decades, some of the MOD members of the Alliance have hosted their own custom BLAST interfaces (Altschul et al., 1990; e.g., FlyBase Consortium. 1999), which have allowed users to search custom databases related to those model organisms, e.g., subsets of related species or molecular clones and display BLAST hits in Genome Browsers aligned with current gene models. We are now developing an updated and integrated Alliance BLAST that optimizes sequence analysis across model organisms, and we have begun to update BLAST at individual MODs. The new WormBase BLAST is now available online, and simultaneously, the FlyBase BLAST system has been replaced and is currently online, with management facilitated through configuration files on GitHub.

The Alliance BLAST will significantly improve the user experience. We envision that BLAST systems, currently powered by SequenceServer (Priyam et al. 2019), will deliver an integrated interface by linking results to Genome Browsers and Alliance gene pages (**Figure 4**). This tight connection allows users to navigate seamlessly between their BLAST results and the wealth of information available within the Alliance, enhancing the efficiency and depth of genetic research. For example, users can retrieve BLAST results for a sequence of interest and then easily navigate across Genome Browsers for different organisms, with a comparison to different tracks revealing how that sequence aligns with gene models, variants, and experimental tools (**Figure 5**). From a project perspective, developing Alliance BLAST with a common cloud-optimized infrastructure will increase efficiency by reducing the cost of compute overhead and eliminating the need to manage separate MOD systems, which will then allow more focus on developing new functionality to support researchers. Our focus in the upcoming year is directed toward enhancing the user interface, reflecting our commitment to providing an intuitive platform for researchers in model organism genetics. We plan to produce more analysis tools as part of the evolving Alliance portal, thereby broadening the range of resources available for genetic research within the community.

**Figure 4.**
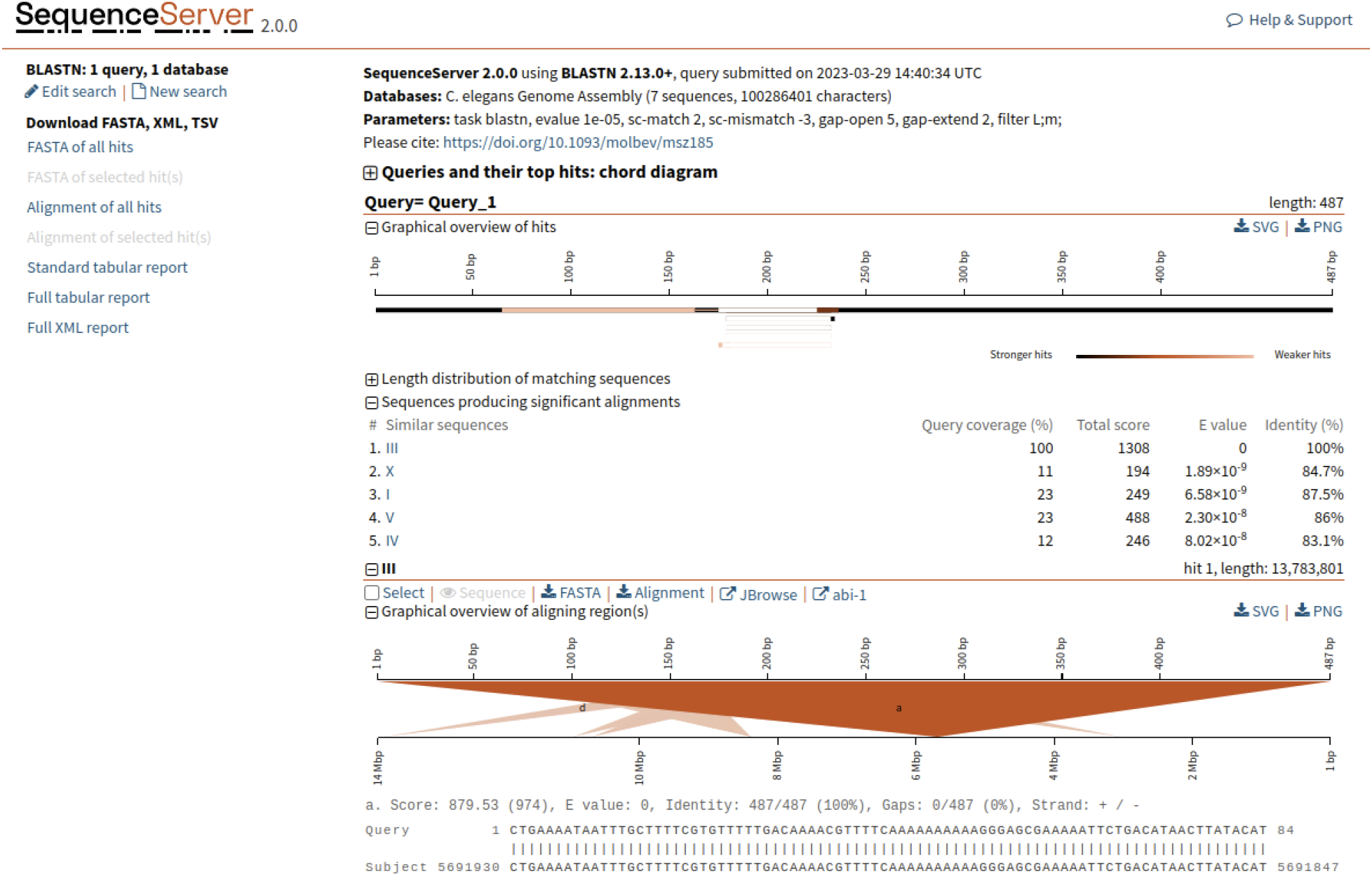
Screenshot of results from the Alliance SequenceServer BLAST tool. The results have been enhanced relative to the default Sequence Server results page by the addition of links to Alliance JBrowse and to the corresponding gene page (in this case, *C. elegans* abi-1) at the Alliance website for each BLAST hit.

**Figure 5.**
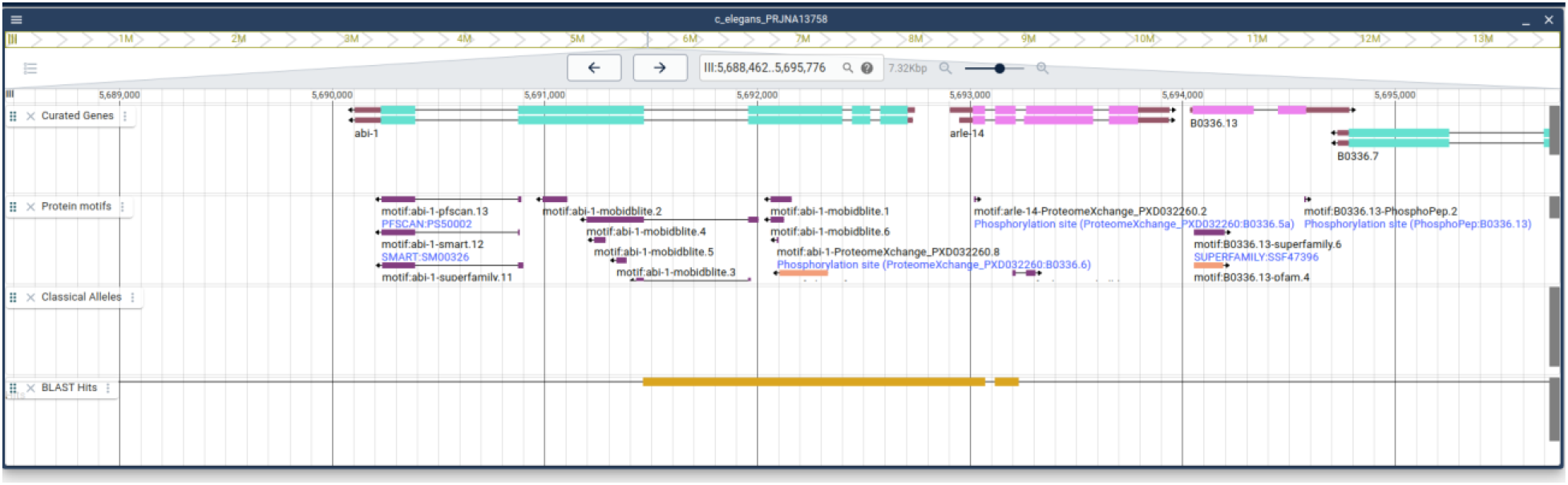
Output of a BLAST search. After a user clicks on the JBrowse link for a BLAST hit they are directed to the web service where they will see a track for the BLAST hit and how the hit aligns with other tracks.

## AllianceMine

AllianceMine, a sophisticated, multifaceted search and retrieval tool that utilizes the InterMine software (Smith et al., 2012), offers a unified view of harmonized data, enabling advanced queries across multiple species. For instance, gene lists can be processed as input and simultaneously query different annotations, such as ’Show me genes associated with a (specific disease term)’ (**Figure 6**). The results from queries can be combined for further analysis, and saved or downloaded in customizable file formats. Queries themselves can be customized by modifying predefined templates or by creating new templates to access a combination of specific data types. Thus, this powerful tool can be used in multiple ways - for search, discovery, curation, and analysis.

**Figure 6.**
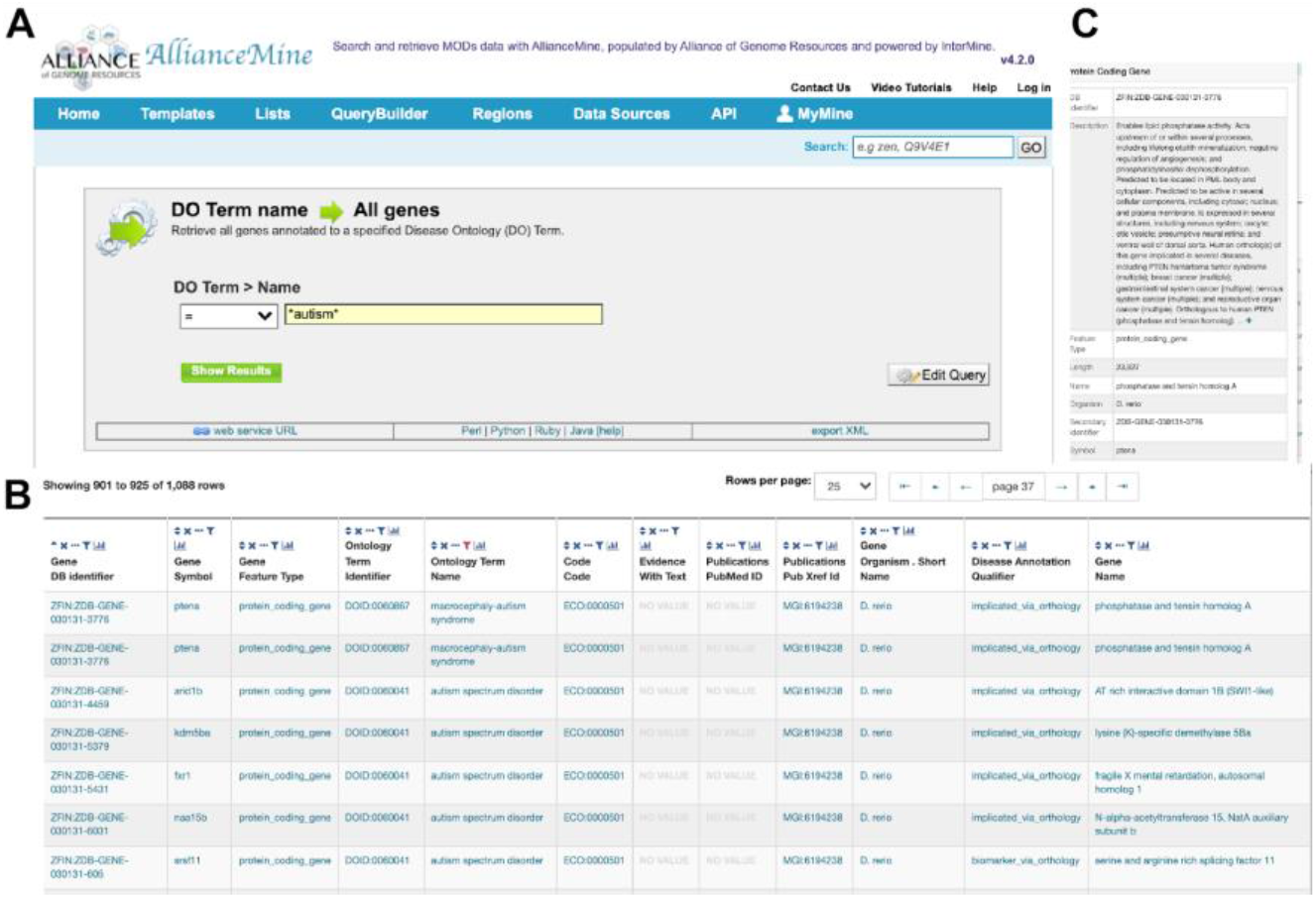
AllianceMine example. Using a simple template, a disease ontology (DO) term is chosen, and all genes associated with this DO term are returned in a downloadable table.

AllianceMine currently showcases harmonized data encompassing genes, diseases, Gene Ontology (GO), orthology, expression, alleles, variants, and FASTA formatted genome sequences. The tool also offers predefined queries or “templates” for cross-species searching. Continual optimization will ensure timely data synchronization with the main Alliance site, as well as integration of newly harmonized data types. Another aspect of improvement will be the addition of more templates, widgets, and pre-compiled lists, which can serve as logical input for templated queries.

## SimpleMine

We designed SimpleMine for biologists to get essential information for a list of genes without any command-line or programming skill, or patience to learn the awesome power of AllianceMine discussed above. Users can submit a list of gene names or IDs to access more than 20 types of essential data with which they are associated. The results are one line per gene with detailed information separated by four types of separators: tab, comma, bar, and semicolon. Users can choose to display the output as HTML or to download a tab-delimited file. Alliance SimpleMine contains ten species curated by the Alliance MODs. It provides easy gene name/ID conversion among MOD ID, public name, NCBI, PANTHER, Ensembl, and UniProtKB. Users can find summarized anatomic and temporal expression patterns, variants, genetic and physical interactions. Other essential gene information includes disease association and orthologs among all ten species. The infrastructure of SimpleMine allows users to perform species-specific searches for lists of genes that take about two seconds to return results, or mixed-species searches that take about 10 seconds to complete.

## Pathway displays with metabolites (GO Causal Activity Models; GO-CAMs)

We have implemented a pathway display on Alliance gene pages, which presents both GO-CAM (Thomas et al., 2019) and Reactome pathway (Milacic et al. 2024) models. The display queries both the Reactome and GO APIs, and shows the number of pathways from each resource that contain the gene of interest. If a gene appears in multiple pathways, users can select which pathway to display. For the GO-CAM models, the viewer has been improved relative to previous releases of the Alliance website (**Figure 7**). First, the layout has been improved to show clearly the overall causal flow through a pathway, from top to bottom and branching as necessary. Second, the viewer displays not only the activities of genes/proteins in a pathway, but also metabolites, which is particularly useful for visualizing metabolic pathways. These metabolites may be either intermediates in a pathway, or regulators of a protein activity. For signaling pathways, we distinguish between direct and indirect regulation, and between positive, negative, or unknown effects.

**Figure 7.**
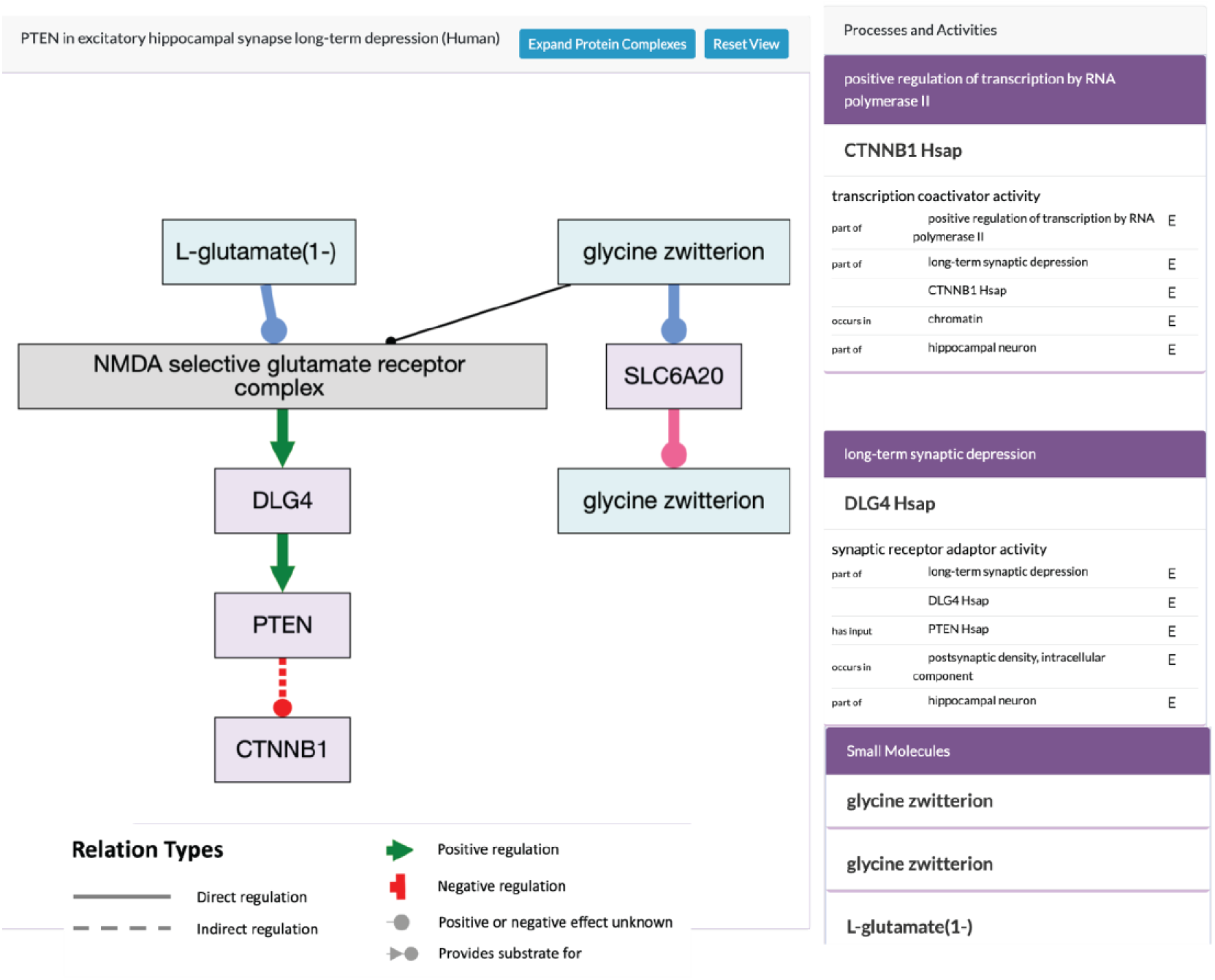
Alliance Pathway Viewer. The pathway widget displays gene products (light purple rectangles), protein complexes (light grey rectangles) and chemicals (light blue rectangles) and the flow of information and material between them (relations). These relations, shown in legend indicate direct or indirect regulation that can be positive, negative or of unknown effect direction.

## Harmonized Data Models

A key requirement of the data transition from individual MODs to the Alliance infrastructure is collective data harmonization so that existing analogous MOD data classes (types/tables) can be loaded into Alliance databases using a consistent data schema and language for communicating about such data classes. A first step in this process, curators from each Alliance knowledge center communicate about their respective existing data classes with an aim of agreeing on which data classes are analogous and therefore should be treated as a single, consolidated data class in the Alliance infrastructure. Next, curators need to align the properties (table columns) of the consolidated data class to agree on property identity alignment and basic data structure including whether these properties are required and/or defining, what values should be stored for these properties, and whether these entity-property-value associations/triples require their own respective metadata and/or evidence records. We use a data modeling language, the Linked Data Modeling Language (LinkML) for these purposes.

Over the last two years, Alliance LinkML modelers have converged on common data modeling patterns that can be reused for each class and property based on the nature of each class property, enabling a standard workflow and implementation to be followed in each case. The LinkML specifications, authored in human-readable YAML files, are used to (programmatically) generate JSON schema specifications that Data Quartermasters (DQMs) can use to generate and validate data files to be submitted to the persistent store. These specifications also inform curation software developers how to generate initial backend (Java models and APIs) and front end infrastructure (curation user interface data tables and detail pages) to be populated once such code is deployed to the production environment of the curation tool and DQMs are ready with their data files. Once DQMs have submitted their data files for a particular data class, the data are loaded into the persistent store with a number of validation and reporting steps (see persistent store architecture description below) and should automatically be populated into the respective data tables and detail pages in the curation interface. The data, having been harmonized, ingested, validated, and displayed to curators in the curation software, can now flow through to the public site according to the data pipeline described (see persistent store architecture description below).

Many Alliance data classes have completely (or nearly completely) harmonized data models in LinkML (see https://github.com/alliance-genome/agr_curation_schema) including: disease annotations, alleles, variants, expression annotations, and references. Although many other data classes have partially harmonized models, ongoing and future harmonization efforts will focus on completing harmonized models for the remaining curated data classes: genes, transcripts, proteins, non-transcribed genome features, affected genomic models (AGMs; strains, genotypes, fish), phenotype annotations, molecular and genetic interactions, gene regulation annotations, high-throughput expression dataset metadata (including for RNA-Seq, single-cell RNA-Seq, and proteomics datasets), species, reagents such as DNA clones and antibodies, images, persons, laboratories, companies, and various entity set classes like gene sets, which can be used for storing assay results and performing downstream analyses like ontology term enrichment, alignments, and other entity set processing calculations.

## Persistent Store architecture

We have designed a powerful database system that can handle most of the demands of our project including curation of data, analysis of data and use and display of the data (**Figure 8**). Specifically, we have instantiated a Postgres persistent store database for long-term and persistent storage of Alliance curated data contributed by Alliance member databases. In parallel to the existing (drop-and-reload) data pipeline (Alliance 2022), DQMs from each MOD now submit data according to our new LinkML schema in JSON format directly to the persistent store for ingestion, validation, and curation via create-read-update-delete (CRUD) operations enabled by a curation API library and Prime React user interface (UI). A data pipeline has been established to provide data from the persistent store Postgres database to our Alliance public website APIs and front end web user interfaces and to other tools and services.

**Figure 8.**
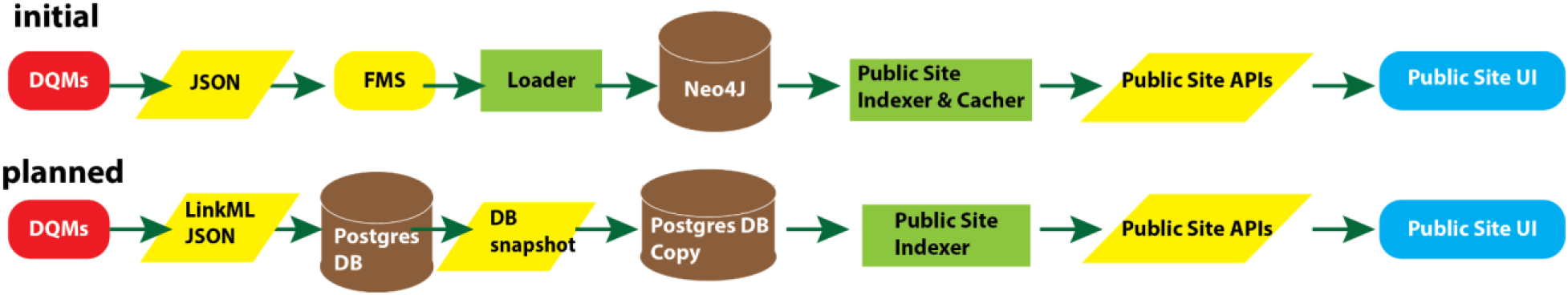
Evolution of Data Flow. Graphical summary showing the design of short term infrastructure initially deployed to support rapid delivery of unified data to the community and the planned production system. Red, data quartermasters at MODs; Yellow, data; Brown, database; Green, transformations; Blue, user interface.

LinkML-based JSON files are ingested into Postgres with validation to ensure: (1) recognition of submitted entities such as genes, alleles, affected genomic models (AGMs; e.g., strains, genotypes), publications, experimental conditions, and ontology terms, (2) recognition of references to such entities in annotations and associations, (3) no entry of duplicate entities, and (4) proper handling of obsolete entities. Every file load is accompanied by a report (in Postgres and the curation UI) indicating (1) the recognized MD5 sum and size of the (uncompressed) file submitted, (2) the success or failure of the load, (3) the number of entities recognized in the submitted file, (4) the number of distinct entities loaded into Postgres, (5) the number and identity of entities (if any) that failed to load and the reason for the failure, (6) a link to download the submitted file, (7) the corresponding compatible LinkML model/schema version, and (8) the MOD data release version corresponding to the data in the file submitted. All of this information can be used by DQMs, curators, and developers to keep track of the fidelity of the data transfer and troubleshoot any issues that arise. Ontology (and other external resource) loads are updated nightly via a cron job to ensure that the latest versions of such data are current. Because the source of truth for MOD data will be transitioned over to the Alliance infrastructure in phases, beginning with a few data types from a few MODs and expanding over time to eventually include all (relevant) data types from all participating MODs, particular logistics need to be addressed. These include recognizing that any discrepancies between data previously submitted by a MOD and data newly submitted from the MOD need to be cleaned up programmatically by removing entities in the database not also submitted in the latest file submission.

To enable create-read-update-delete (CRUD) operations on persistent store data, curation APIs and a curation user interface accessible with Okta authentication have been implemented (**Figure 9**). Curators can now access data tables for the following data types: genes, alleles, variants, affected genomic models (AGMs; e.g. strains, genotypes), publications (accessed via Alliance Bibliographic Central (ABC) APIs), experimental conditions, constructs, disease annotations, molecules (not already managed by Chemical Entities of Biological Interest (ChEBI)), ontology terms, and controlled vocabularies and their terms. CRUD operations have been fully enabled for disease annotations, experimental conditions, and controlled vocabularies, read-update operations have been enabled for alleles and variants, and read operations are enabled for the remaining data types. Work is underway to fully enable CRUD operations on all remaining data classes and their attributes including new data tables for transcripts, proteins, other (non-gene) genome features, expression annotations, phenotype annotations, molecular interactions, genetic interactions, gene regulation annotations, antibodies, images, and more. In addition to data tables presenting all entries of a particular data class, the curation tool also has individual entity detail pages (for example, see an allele detail page https://curation.alliancegenome.org/#/allele/MGI:6446761) for data entry and editing on a dedicated web page for one particular entity. The curation tool also enables user-specific and MOD-specific custom user settings and preferences to provide a user interface most compatible with individual curators’ workflows.

**Figure 9.**
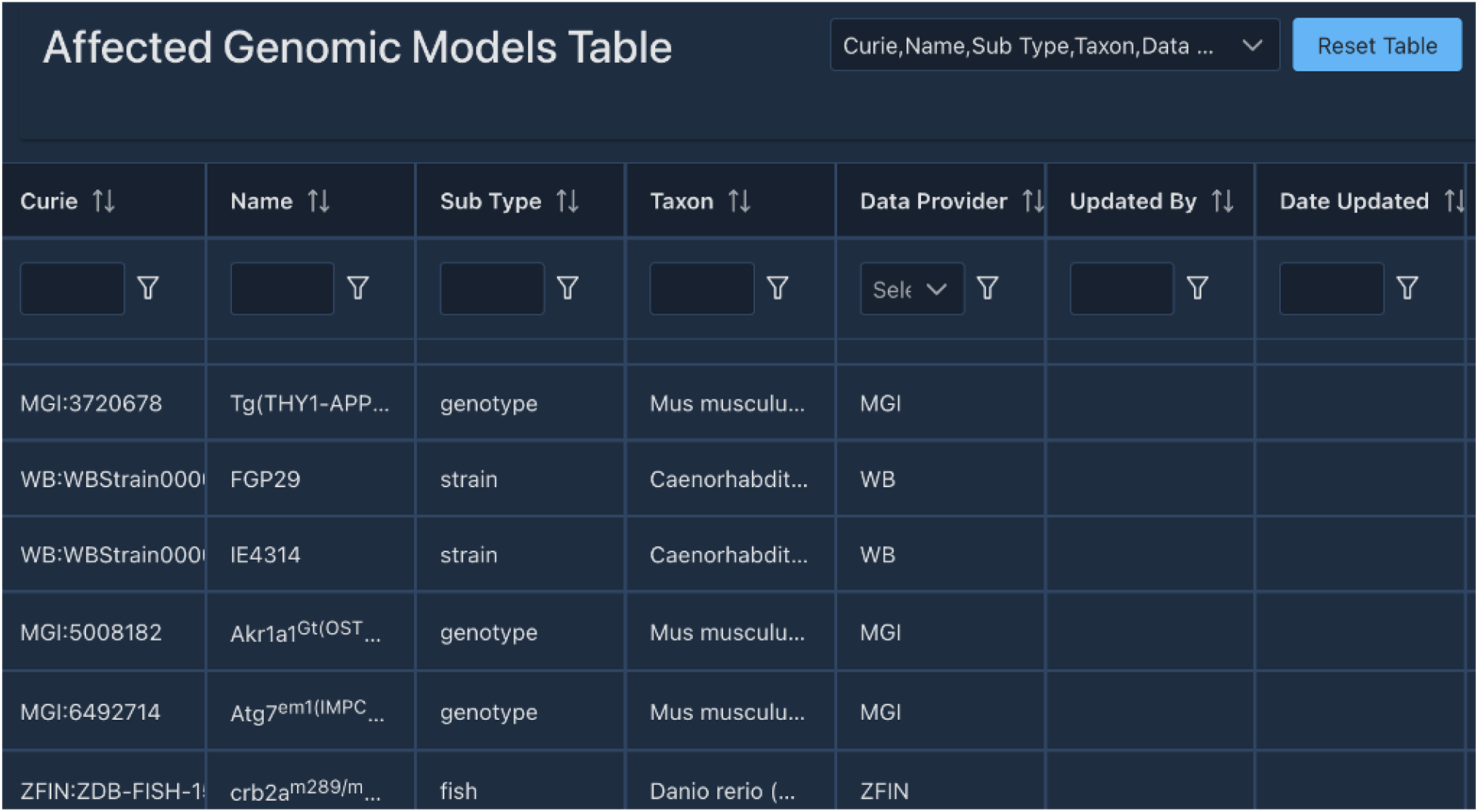
Screenshot of the Alliance curation tool interface showing an example of curated annotations of Affected Genomic Models managed in the persistent store.

Future development plans for the curation tool include: batch creation of data entities (e.g., annotations, reagents), batch editing, data history inspection and auditing, undo and review of latest changes, publication constraints (constrain data view and entry to publication currently being curated), customizations and MOD default settings for new entity creation and detail pages, incorporation of data entity and topic tagging information from the ABC literature store, and incorporation of AI/ML into the curation workflow.

For releases of persistent store data to the Alliance public website, Postgres database snapshots are taken and sent to a separate Postgres instance that feeds the data via the curation APIs (instantiated as a library) into the public site indexer where various data filtering and transformations occur before making those processed data available to our public website APIs via our Elasticsearch index. The Alliance public website user interface, using existing UI infrastructure, is then modified or created to accommodate the data now flowing from the persistent store database.

## Security, stability and backups

All services and data provided by the Alliance to its community are hosted on Amazon web services (AWS). This provides us with industry leading availability of up to 99.99% on services like EC2, which we use to host our virtual servers. We use additional AWS-managed services such as Elastic Beanstalk for application deployment, RDS for hosting our relational (postgres) databases, and Amazon OpenSearch Service for hosting our search indexes, which all provide automatic updates and maintenance for increased reliability. All files hosted at the Alliance of Genome Resources are stored in S3 buckets, which ensures industry leading durability and availability. Furthermore, we make daily backups of our relational databases and have processes in place that enable easy restore of those backups in case of failure or data corruption. All Search indexes are derived from the persistent relational database and can be regenerated at any moment when required.

We make use of separated AWS VPC subnets between public-facing and private systems, and only services requiring public access are given public IP addresses. This ensures that public-facing services such as our curation interface can be accessed by our curators world-wide (through Okta Authentication), although the supporting back-end services such as the supporting databases can be kept private and can be accessed only by authorized internal users by connecting to our internal network through the AWS VPN. Access to all services is furthermore restricted to allow access only to the required ports and services through the use of AWS Security Groups to control the allowed network traffic. AWS IAM users, groups, and roles are used to control the allowed AWS operations and access among Alliance developers. In all cases, the principle of least privilege is applied, so that the potential attack surface is reduced to a minimum (for example by not granting blanket AWS admin permissions to developers who do not have an AWS admin function). Access keys to any system can be revoked when misuse of those access keys is detected. Furthermore we configured our github repositories to be scanned automatically for accidental secret credential leakages through the use of GitGuardian.

## Literature Acquisition

We designed and are implementing a literature system, Alliance Bibliographic Central (ABC), that will support curation, and in the future, end users. The ABC supports the tasks of reference acquisition, triage, and curation workflow management. Specifically, the ABC is an ecosystem of online tools and supporting Alliance databases that manage all references and related metadata that are ‘in corpus’ for the member MODs.

During the past year, we focused on literature acquisition. Literature acquisition at the Alliance begins with automated, organism-specific PubMed queries to retrieve candidate references for each MOD’s corpus. References matching the search criteria are then added to the ABC by assigning an Alliance reference id and importing associated bibliographic information to the database. Subsequently, curators manually sort references as either ‘in’ or ‘out of corpus’ based on the curation policies of the MOD and eliminate any false positive results from the initial search. Once references are sorted, they enter MOD-specific curation workflows supported by task-specific ABC curator interfaces to, for example, add reference files, manually tag references with specific entities (e.g., genes, alleles, and data types) and topics (e.g., phenotypes, anatomic expression) using the Alliance Tags for Papers (ATP) ontology, and merge duplicate references. In addition to adding reference files manually, the full text of ‘in corpus’ references included in the PubMed Central (PMC) open access set is also automatically downloaded. Curators may also use the ABC to add non-PubMed references. An additional key feature of the ABC is a search interface that allows curators to retrieve references based on various criteria including their in/out of corpus status, bibliographic data, and publication data range, if desired. Reference acquisition functionality can easily be extended to integrate additional MODs into the Alliance infrastructure.

To facilitate reference data exchange between the Alliance and MOD databases, the MODs provide a mapping file that associates MOD reference CURIEs (Compact Uniform Resource Identifier) with PMIDs, e.g., ZFIN:ZDB-PUB-181026-2 - PMID:30352852. The MODs also provide reference CURIEs and data for references not included in PubMed but used by the MOD, such as internal curation references and those published in a journal not yet indexed at PubMed.

Over the past 25-30 years, Alliance member databases have independently developed methods to acquire, triage, and curate their respective literatures. Having implemented a common literature curation interface, database, and full text acquisition system, the ABC is now poised to expand its functionality by incorporating ML methods developed by, and in production for, a subset of Alliance members to all groups. For example, automated pipelines that recognize entities (e.g., genes, alleles, strains) as well as data types (e.g., phenotype, genetic interactions) can be developed for new groups with results stored centrally in the Alliance literature database. Incorporating more automated methods will allow faster association of the published literature with relevant biological concepts, information that can be displayed on future Alliance references pages while the papers await detailed full curation. Centralized literature infrastructure will also support other curation pipelines, such as community curation by authors, which can then be more readily implemented for additional Alliance member communities thus providing another avenue by which curated data can be swiftly included in the Alliance. Lastly, the common literature tool will allow Alliance biocurators to coordinate curation of multi-species references that will provide users a facile way to find and view cross-species research exploiting the strengths of each Alliance model organism, a primary goal of the Alliance.

## Textpresso

Textpresso is a full-text literature search engine that gets power from its single-sentence scope, focus on a specific model organism (or topic), and categories of semantically or biologically related terms (**Figure 10**; Müller et al., 2004; Müller et al. 2018). It has been used extensively by WormBase and SGD curators, as well as *C. elegans* and *S. cerevisiae* researchers in addition to other MODs (Van Auken et al., 2012; Bowes et al., 2013)

**Figure 10.**
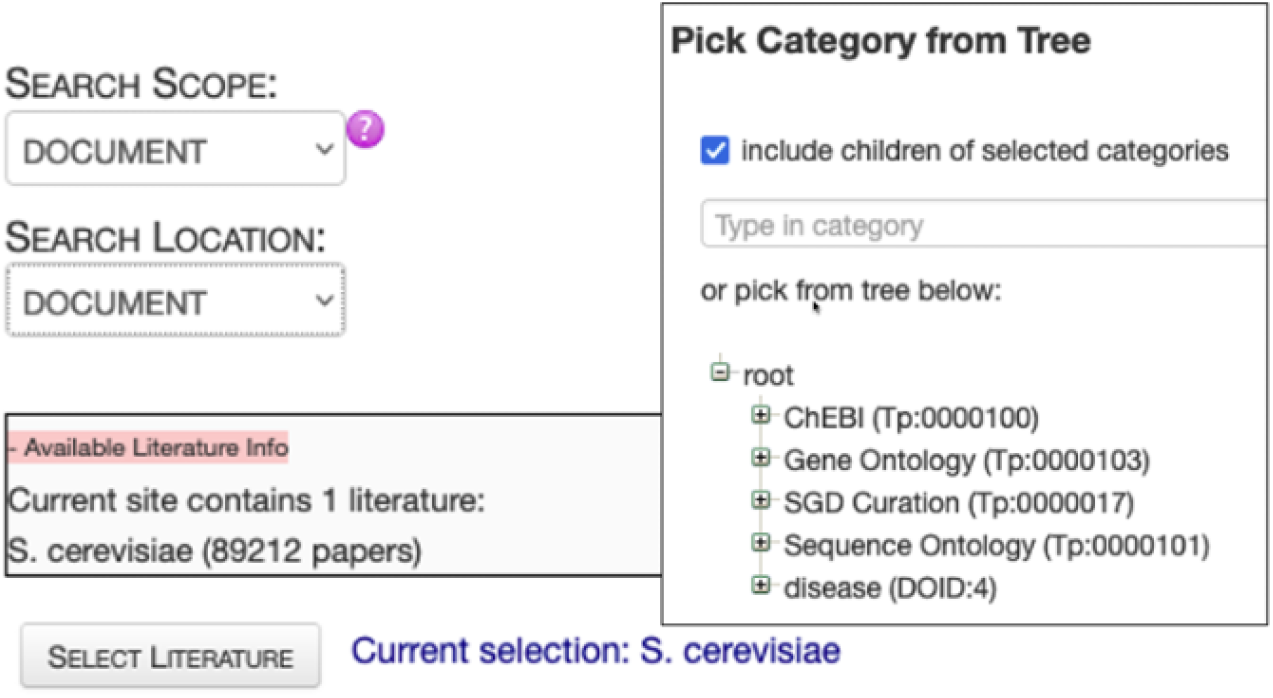
Textpresso for SGD literature at the Alliance. (http://sgd-textpresso.alliancegenome.org/tpc/search)

The Alliance is committed to creating Textpresso instances tailored to the unique needs of each member database, all of which will be managed within the Alliance software ecosystem and connected to the ABC as a single reference data source. This will reduce the overhead of managing Textpresso at individual MODs while also simplifying development and deployment of new features. Users will benefit from simplified access to Textpresso from the Alliance website. We also plan to integrate Textpresso searches further into specific Alliance web pages such as gene or allele pages. Users will be able to obtain additional references to biological entities through Textpresso searches, adding information from potentially non-curated literature to the list of curated references currently linked on those pages. Textpresso will be available to Alliance biocurators and to the general public through MOD-customized websites and via APIs for programmatic access.

## Artificial Intelligence (AI)

The Alliance member MODs have a track record of implementing ML tools to enhance triage and curation efficiency. Notable examples include RGD’s early adoption of UIMA standards and the development of the OntoMate system (Liu et al. 2015), as well as WormBase’s creation of Textpresso (Mueller et al. 2004) and document classifiers for paper triage.

The rise of Large Language Models (LLMs), like BERT, short for Bidirectional Encoder Representations from Transformers, and ChatGPT, has transformed the NLP landscape, but questions about their accuracy and “hallucinations” remain. The Alliance aims to harness LLMs for tasks such as document classification, Named Entity Recognition (NER), sentence classification, assisted-triage and curation and to build a natural language query system to simplify access to its underlying structured data.

In the realm of AI/ML, Alliance members have developed classifiers for determining with high accuracy whether papers returned from automated PubMed queries should be kept in their corpus or discarded. The Alliance is developing a central solution by providing this type of in/out corpus classifier to all members.

Efforts are also underway to improve existing species-specific entity extraction and classification models, with a focus on incorporating human feedback in the loop and continuously training models based on data validated by professional biocurators and community curators. A centralized interface for “topic and entity tag” addition and validation during triage and curation is under development as part of the ABC. The interface allows curators to associate tags with publications and at the same time validate (or invalidate) results extracted from AI/ML methods. This interface will streamline the collection of valuable training and testing sets and will allow a more systematic approach to the creation and comparison of different AI/ML models. Future plans include development of tools for creating training sets and a model manager for tracking ML models’ performance. Integration with specialized biocuration tools such as Ontomate and Textpresso is part of the strategy, with a vision of harmonizing AI/ML solutions across member sites.

Furthermore, the Alliance is adopting Evidence and Conclusion Ontology (ECO) terms to record systematically the type of evidence, e.g. neural network method evidence, and assertion method, e.g. automatic assertion, used for reference flagging and triage. This is especially relevant for topic and entity tags. Using ECO terms aligns with FAIR data principles and offers transparency in curation workflows.

We will also explore the use of AI/ML in gene function summarization. Included on gene pages at the Alliance are short textual gene summaries based on curated and structured data that provide users a quick overview of gene function. The current automated system for generating gene summaries has produced more than 160,000 summaries (Alliance version 6.0.0) for nine species, including humans (Kishore et al., 2020). However, to increase the coverage of genes further, we will explore the use of LLMs. This is especially relevant for less-studied genes with few curated, structured data, and for scaling and upkeep of the summaries to match the rate of new gene data from publications. We will use prompt engineering and finetuning of LLMs to improve accuracy of the generated summaries. As part of a continual improvement process, we will ask professional biocurators to evaluate summaries, and we will develop a scoring system based on several features such as readability of summaries, inclusion of key gene data, and checking for inaccurate and false data. To improve and keep gene summaries up to date, we plan to retrieve newly published articles that contain gene data that were not available when the LLM was trained and add extracted relevant text from the identified articles to the LLM prompt. To do so, we will use tools such as Textpresso (Muller et al., 2004) and Ontomate (Liu et al., 2015)

## Application Programming Interfaces (APIs)

Application Programming Interfaces (APIs) are a key component of Alliance Central’s data services infrastructure for rapid, modular software development. We currently support a dozen APIs with hundreds of endpoints (**Figures 11**, **12**). New APIs will be added as data harmonization and modeling of additional data entities are completed. We will expand public site APIs to generate all data needed for SimpleMine, AllianceMine, etc. from single endpoints.

**Figure 11.**
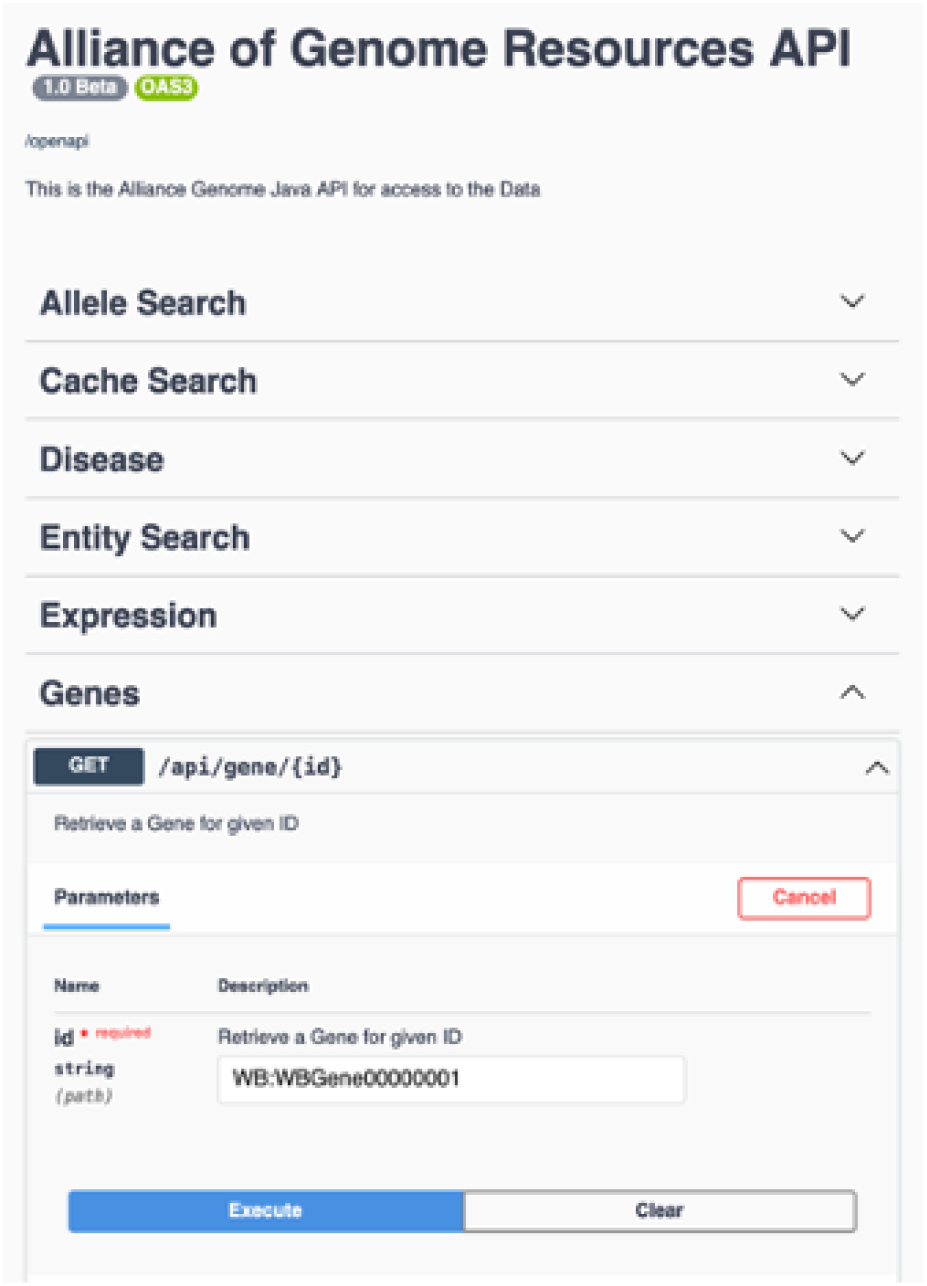
Swagger interface for the Alliance APIs.

**FIgure 12.**
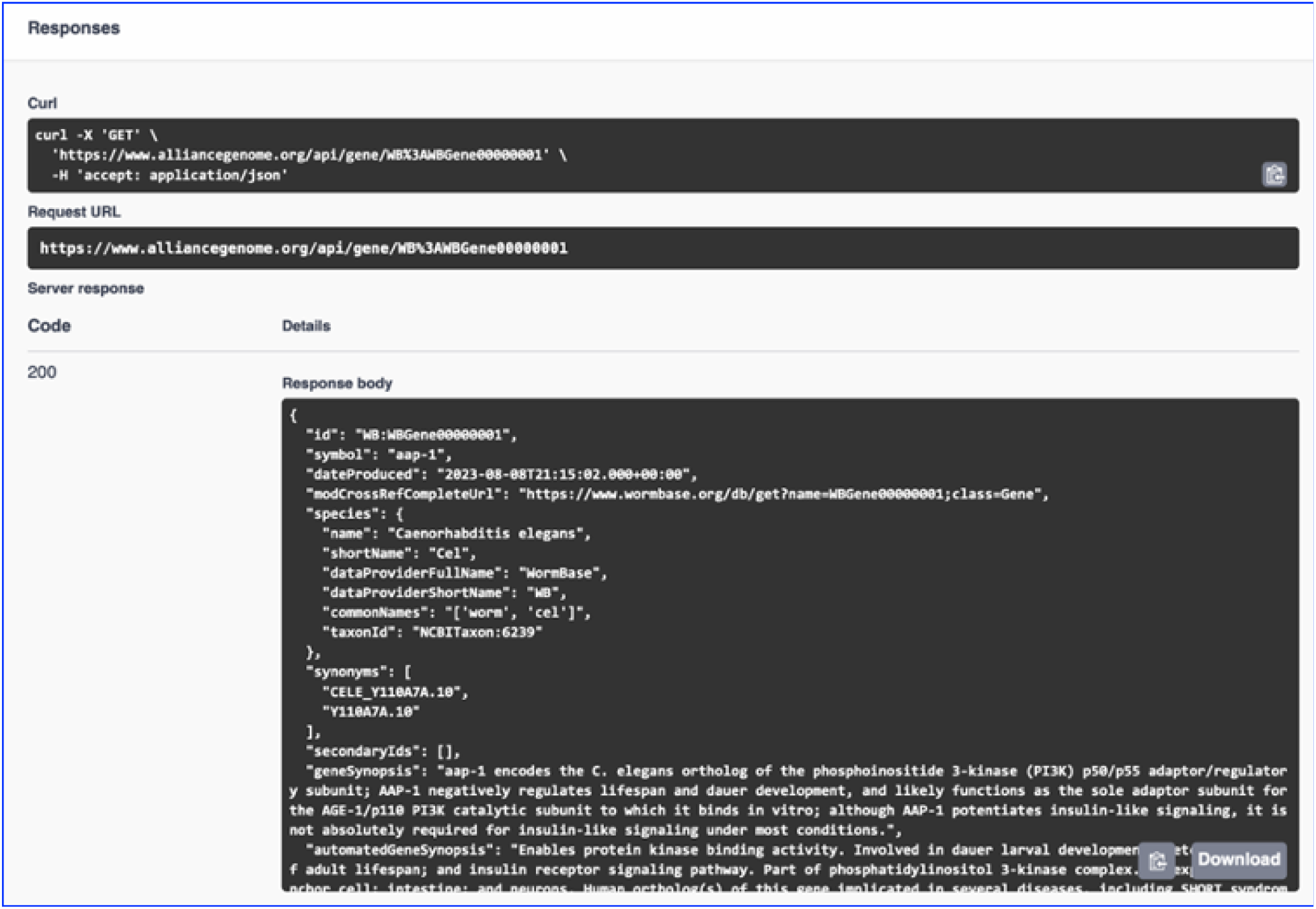
Example of API output.

Current APIs include Public site APIs (agr_java_software in the GitHub repo) and APIs available from a public Swagger UI page. Because the public APIs support only GET endpoints, they do not require authentication. All APIs that support both GET and PUT/POST/DELETE endpoints do require authentication. Some of the key API endpoints available at https://www.alliancegenome.org/swagger-ui/ are: gene-summary, gene-disease, gene-interactions, homologs-species, allele-phenotypes, expression ribbon-summary, etc.

## Data preservation in external repositories

The Alliance of Genome Resources is committed to the long-term preservation of digital objects (annotations) and resources (e.g., ontologies and software) that are central to the management and integration of functional knowledge about the genomes of diverse model organisms. As part of this commitment, the annotations and resources generated by Alliance members are integrated into many long-standing external public bioinformatic resources (e.g., Ensembl, UniProt, NCBI). Distribution of Alliance annotations from multiple sources provides a degree of redundancy that contributes to data stability and preservation. Alliance maintained ontologies and annotations and are also deposited into third party repositories that fulfill Open Science principles (see below). Leveraging community repositories ensures the data products and resources remain accessible to the research community even if the Alliance and/or its members cease operations.

Ontologies that Alliance members maintain are also available from long-term repositories including the OBO Foundry (https://obofoundry.org/) and Zenodo (zenodo.org). Annotations related to gene expression, function, phenotype, disease associations, etc. that are generated by Alliance members and are available on the Alliance Data Downloads page are archived in Zenodo. Software developed as part of the Alliance of Genome Resources knowledge commons platform is available from GitHub (https://github.com/alliance-genome). The external repositories used by the Alliance of Genome Resources include the *OBO Foundry* that was established in the early 2000s as a community-based initiative for development and maintenance of biological and biomedical ontologies using standardized practices. The Foundry is the ontology repository of choice for the Alliance because it is widely recognized as an authoritative source of well-maintained ontologies for biology and biomedical research.

*Zenodo* is a general purpose repository maintained by CERN (European Council for Nuclear Research) for storing and sharing documents, data, and other digital research materials across many disciplines. Zenodo is a repository of choice for the Alliance, in part, because of the commitment by the European Commission to support Zenodo as long as CERN exists.

## Disease Portal(s)

Providing users with ready and easy access to curated and harmonized model organism disease data and tools is crucial to accelerate research related to the pathogenesis of human disease. The Alliance has a wealth of disease-relevant data from eight model organism species and human data, such as: genes, alleles and variants implicated in disease, genotypes and strains that serve as disease models, and related data such as modifiers (herbals, chemicals, small molecules, etc.) that ameliorate or exacerbate the disease condition and may serve as candidates for potential drug development. To provide an easy entry point for clinical researchers and human geneticists to access the consolidated data and tools, we are in the process of designing and implementing a topic-specific resource--an Alzheimer’s disease (AD) portal that will serve as a paradigm for other disease portals (**Figure 13**). The AD portal will include: orthologous genes in animal model systems, models with a mutation orthologous to one in a patient group, models with a specific set of phenotypes, and/or modifiers that have been shown to alter the disease condition. Building on the experience and pages developed for the AD portal, we will expand this paradigm to other disease portals. Features planned for the disease portal with AD as an example include: a home page with an overview of the data in the portal, an autocomplete search box, links to other AD resources, and a list of the most recent papers from PubMed and/or from the ABC store (see example portal page below). The pages in the portal will be modeled on existing pages at the Alliance and will include gene summaries, alleles and variants, phenotypes, gene interactions, pathways, biological processes (based on GO), gene expression, etc. We also plan to provide visualizations of data analysis, for example, diseases that share genes and protein interactions that may point to common underlying molecular mechanisms. Up-to-date data sets, e.g., genes, strains, modifiers (drugs, chemicals, herbals, etc. shown to either ameliorate or exacerbate phenotypes) will be available as downloadable files. Disease-specific data sets will also be available for query from AllianceMine. We will also provide up-to-date links to disease-specific literature, and search capabilities through literature search engines such as the Textpresso instance dedicated to AD (http://alzheimer.textpressocentral.org; corpus size - 96,000 papers).

**Figure 13.**
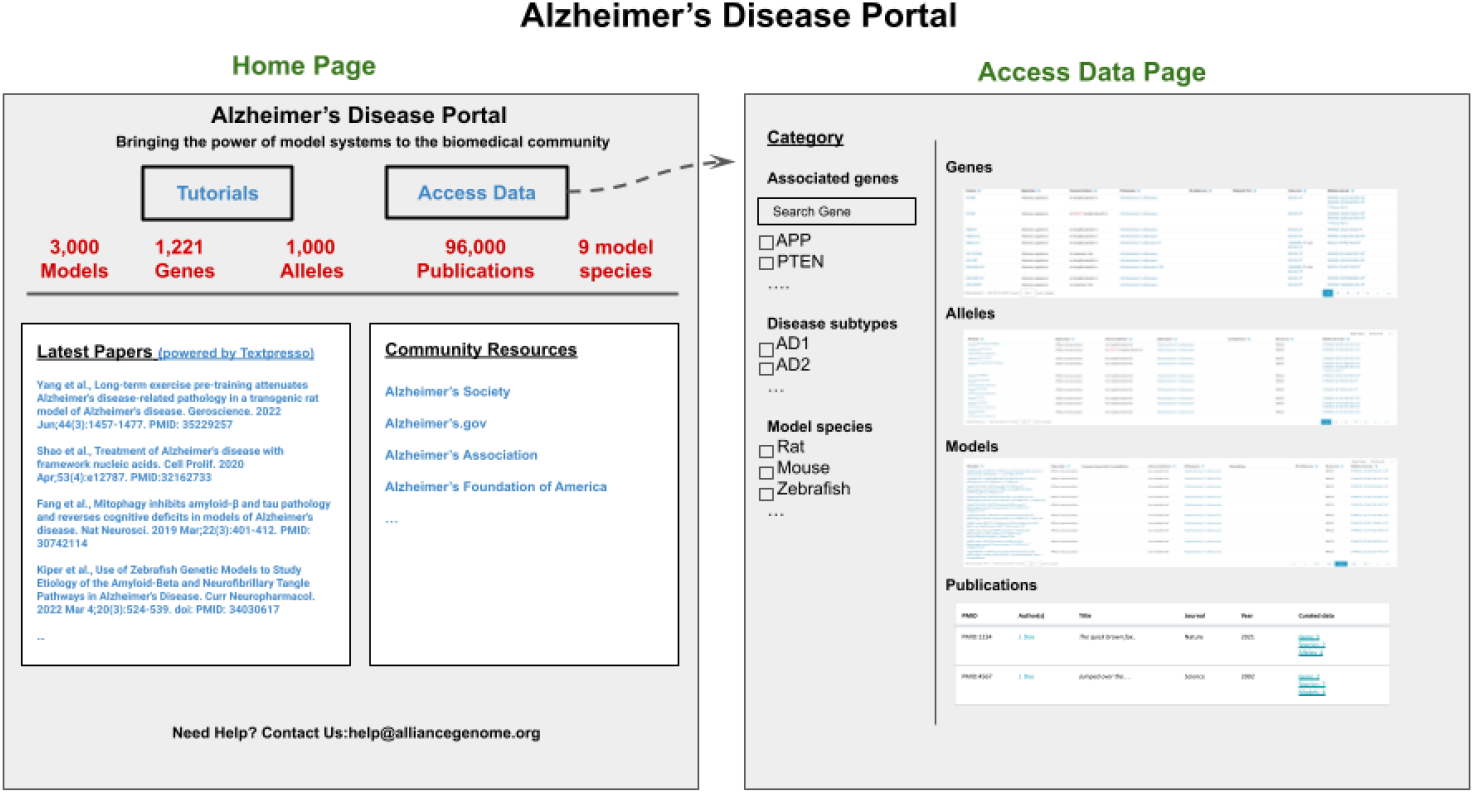
Mockup of the Alzheimer’s Disease Portal showing the Home page and the Data access page. These views illustrate the type of information that will be available with a disease-focus.

## Outreach and interactions

### The Alliance Helpdesk

We established a common help desk email address that is featured prominently on the Alliance website header and footer under “Contact Us”. All inquiries submitted using this email are logged as tickets in the Alliance Jira software system. Members of the User Support Working Group respond to user questions and inquiries in a timely manner, typically within 48 hours. Time to resolve user inquiries depends on the nature of the question or request. The Jira system tracks open tickets, forward tickets, tracks their active/resolved status, and classifies them by subject. We use the information, in part, to evaluate the design and utility of our user interfaces. For example, if particular questions or subjects arise frequently, we re-evaluate the design and wording of the search form and/or results display that caused user confusion.

### Online documentation

We provide extensive user documentation about using the Alliance data resources under the Help menu on the homepage (https://www.alliancegenome.org/help). The online documentation provides guidance on such topics as how to use the search functions, defines acceptable field parameters, and provides explanations of the displayed results. The User Support Working Group also works closely with the User Interface Working Group and the Developers to craft text for tooltips displayed on user interfaces.

### Frequently Asked Question (FAQ) pages

The FAQ/Known Issues page provides answers to commonly asked questions about the Alliance and also describes any known issues associated with a particular software release. The link to the FAQ page is featured prominently on the Alliance home page under the Help menu.

### Illustrated tutorials and videos

We maintain several types of tutorial options that are accessible from the Help menu (https://www.alliancegenome.org/tutorials). The most requested types of tutorials are illustrated guides with screenshots on how to use various features of the Alliance web portal. When new functionality is released, we post to social media channels and issue “Tweetorials”. Short video tutorials are disseminated through the Alliance YouTube channel.

### Alliance User Community Forum

The Alliance supports a centralized community discussion board implemented in Discourse (https://community.alliancegenome.org/categories) (**Figure 14**). Each model organism represented in the Alliance is represented as its own Discourse category with model organism specific threads for news, discussion, and reagent information. The forum also includes categories for job postings, meeting announcements, and general information about the Alliance of Genome Resources. Alliance members with existing on-line community forums are migrating users to the Alliance Central forum.

**Figure 14.**
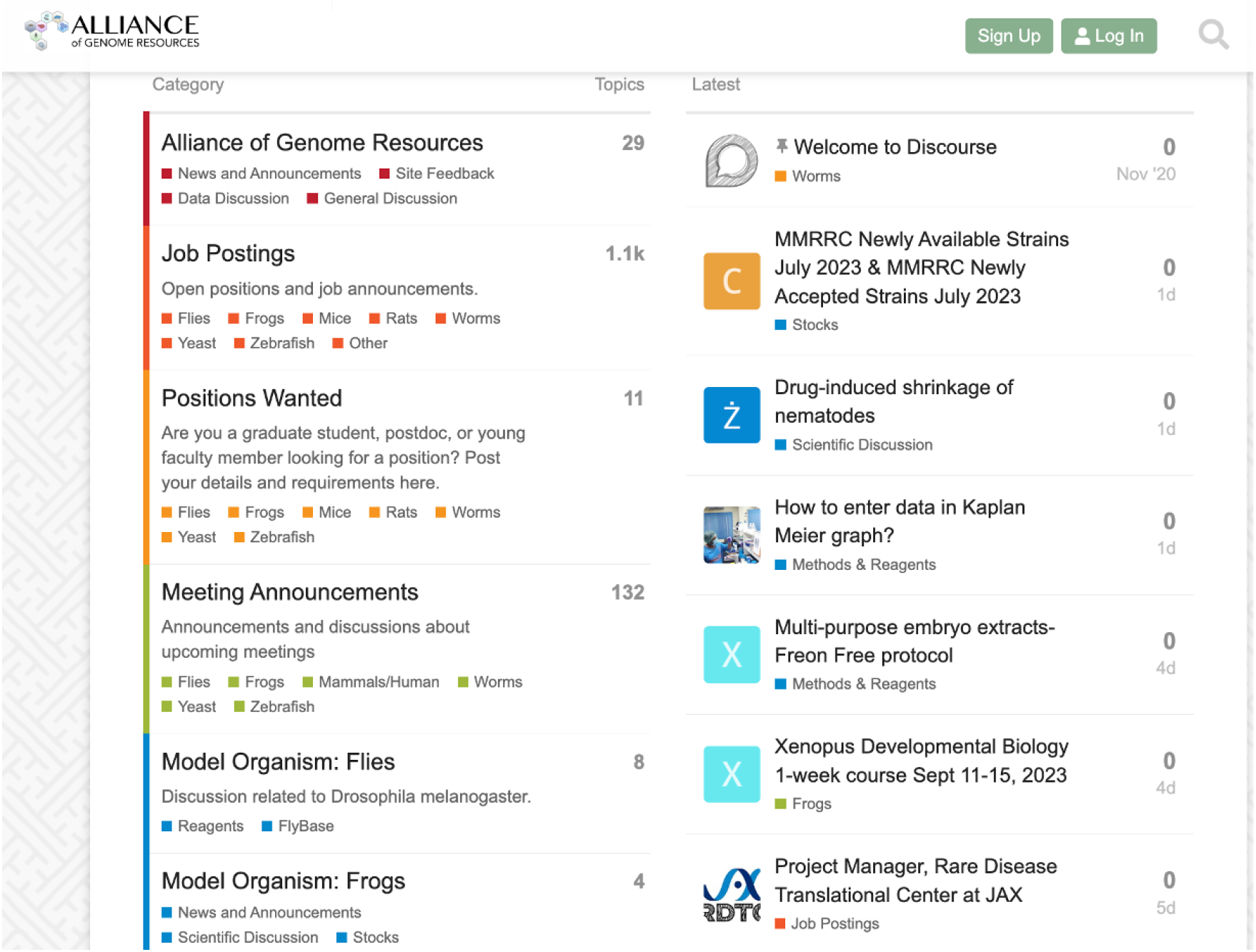
Alliance community forum home page.

Users are not required to register to access the forum but must register to post messages, questions, and announcements. On average, ∼1,000 users a day access the forum. Posts include jobs open and sought, news, meeting announcements and discussion of research approaches, reagents and interpretation.

### Social Media

In addition to a News and Events header that links to software release notes and other Alliance Central updates, the Alliance uses standard social media venues to engage with the user community, including FaceBook (www.facebook.com/alliancegenome/), Twitter (now, X) (twitter.com/alliancegenome), Mastodon (https://genomic.social/@AllianceGenome), and Bluesky (https://bsky.app/profile/alliancegenome.bsky.social).

## Prospects and Challenges

### The long tail of data

One challenge in the central Alliance infrastructure providing support for the union of MOD and GO features is the many unique dataset displays and tools that have evolved in the individual MODs over two decades. Among the 8 resources this comprises 150 years of branch length! Although horizontal tool transfer has occurred, it is not complete. We are taking a few approaches to this problem. In some cases, where the data are stand-alone, we will simply move the data and code to the Alliance. In the short term we will likely run tools off their existing servers. As tools age out, we will evaluate whether there is a broader mandate for that feature, and if so, implement it in the context of the Alliance.

### The tail of not-yet harmonized data

There are types or aspects of our data that can be harmonized but have not yet been so. We adopted LinkML to help with harmonization because it provides a common language to represent structured data. The use of this language has spread to the point where our progress on harmonization is much more rapid.

### AI

As discussed above, we are actively considering AI/ML applications throughout the project. Our practical approach is driven by us being subject matter experts. Because we have relied on human expert curation, we are in a unique position to evaluate and use the output of various AIs.

### Community curation

Some Alliance MODs employ community curation pipelines to engage authors in curation of their papers. For example, FlyBase utilizes the Fast Track Your Paper (FTYP) (Bunt et al 2012; Larkin et al., 2021) tool that allows users to curate scientific papers, identify data types, and associate relevant genes with the reference. Authors using FTYP ensure their papers appear quickly on the FlyBase website, help highlight data needing manual curation, and prioritize their publication for further curation.

Similarly, WormBase developed ACKnowledge (Author Curation to Knowledgebase; Arnaboldi et al., 2020), a semi-automated curation tool that lets authors curate their publications with the help of ML. Authors receive an email with a link to a form pre-populated by document-level classifiers that identify data types and several NER pipelines that extract lists of entities. Authors can correct and validate the extracted data using the form and submit curated information to WormBase. We will continue to provide these services to our community and develop a unified infrastructure which will help expand the service to other member communities.

Several Alliance members also collaborate with publishing groups, such as microPublication Biology (https://www.micropublication.org/) or the Genetics Society of America (https://genetics-gsa.org/publications/), to streamline pre-publication data integrity verification and curation by curators and authors, enabling MODs to quality-check and work with authors to correct data reporting before publication and promptly incorporate it into Alliance Knowledgebases upon article publication.

### Dealing with satellite genomes and genetic models

In addition to the core genomes and associated data, our resources store and present information about the genes and genomes of relatively closely related organisms. For example, WormBase includes some genetically-studied nematodes such as *Caenorhabditis briggsae* that benefit from the rich data models typical of *C. elegans*. Genetic screens and positional cloning (Inoue et al., 2007; Sharanya et al. 2012), CRISPR editing (Cohen and Sternberg, 2019; Cohen et al., 2022; Ivanova and Moss 2023), as well as transcriptomic analyses (Jhaveri et al., 2022) are now routinely done in this species. For the Alliance to take on this responsibility of WormBase, we need to support such satellite model organisms. Our plan is to support community gene structure annotation (e.g., for *Drosophila*, Sargent et al, 2020; for *C. elegans*, Moya et al. 2023) using the Apollo curaton system designed specifically for such activity (Dunn et al., 2019).

## High Throughput expression data and single cell RNA-seq plans

We harmonized high-throughput expression metadata of mouse, rat, yeast, worm, fly, and zebrafish. Users can browse them with species, assay type (microarray, RNA-seq, tiling array, and proteomics), tissue, sex, and curated categories. We plan to add single-cell RNA-seq as a new assay type, making such datasets easily identifiable within our collection, with links to other resources, including Gene Expression Omnibus, EBI single-cell RNAseq Expression Atlas, ans CZI CellxGene, To display the information above, we will implement a content-rich expression detail page that will provide a unified way to access all expression data associated with a specific gene, including link outs to other sources and MOD-specific single-cell RNA-seq gene expression graphs (**Figure 15**).

**Figure 15.**
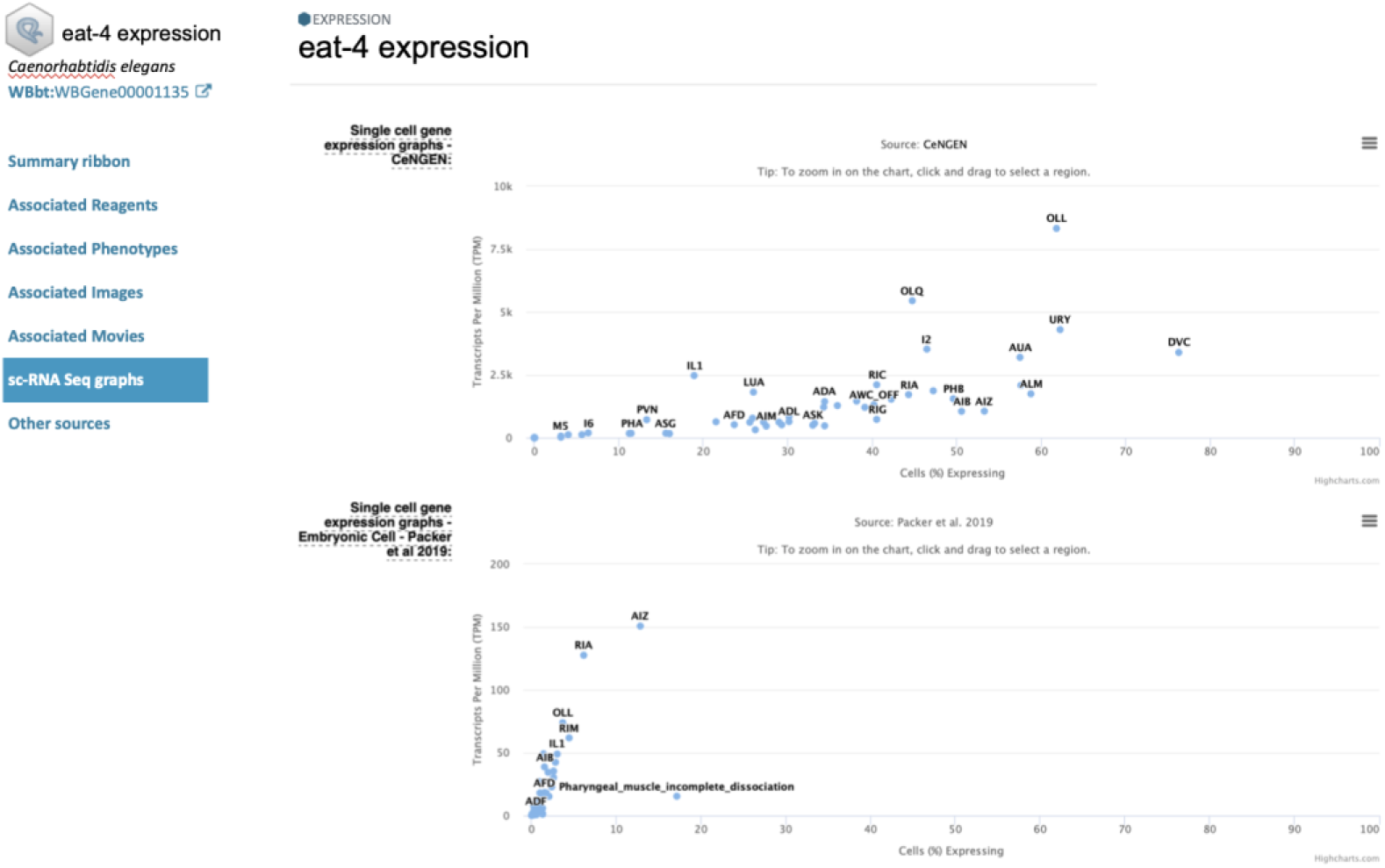
Mockup of an Expression Detail page. This example shows one of the current features of WormBase – single cell data from two studies – displayed on what will be part of an Alliance Gene Expression detail page.

### The Alliance in the ecosystem of knowledgebases

The Alliance has a unique and complementary role relative to other informatics resources that support comparative biology. For example, NCBI’s new Comparative Genomics Resource (CGR) (Bornstein et al 2023) focuses on developing analysis tools and resources for *sequence-based* genome comparisons across a large number of species, the Alliance focuses on standardized annotations, harmonized biological concepts, and comparison of *biological knowledge*. The CGR supports comparative sequence analysis for all eukaryotes whereas the Alliance is primarily focused on model organisms used widely in biomedical research. These model organisms have a tremendous amount of highly valuable genetic, transgenic, and phenotypic data generated with multiple types of assays and are uniquely represented by the Alliance Knowledge Centers. The CGR uses the standardized gene summaries from the Alliance and follows nomenclature and ontology standards developed and maintained by Alliance members. For sequence analysis, the Alliance leverages sequence-based analysis tools developed and maintained by the CGR. Resource developers by and large appreciate the magnitude of the tasks we face in order to provide researchers with the information they need, and strive to fill in the many gaps in services.

## Acknowledgements

We thank our multiple communities for their patience and feedback about the prospect of the Alliance and their love of their own MODs. We also thank the members of our Scientific Advisory Board (Gary Bader, Alex Bateman, Helen Berman, Shawn Burgess, Andrew Chisholm, Phil Hieter, Brian Oliver, Calum Macrae, Titus Brown, Abraham Palmer and Michelle Southard-Smith) for cogent advice, and NHGRI Program Staff (Sandhya Xirasagar, Ajay Pillai, Valentina di Francesco, Sarah Hutchison, and Helen Thompson) for guidance. The core funding for the Alliance is from the National Human Genome Research Institute and the National Heart, Lung and Blood Institute (U24HG010859). The curation of data and their harmonization is supported by National Human Genome Research Institute grants U24HG002659 (ZFIN), U24HG002223 (WormBase), U41HG000739 (FlyBase), U24HG001315 (SGD), U24HG000330 (MGD), P41HD064556 (Xenbase), U24HG011851 (Reactome + GO) and U41HG012212 (GO Consortium), as well as grant R01HL064541 from the National Heart, Lung and Blood Institute (RGD), P41HD062499 from the Eunice Kennedy Shriver National Institute of Child Health and Human Development (GXD), and the Medical Research Council-UK grant MR/L001020/1 (WormBase). Additional effort was supported by DOE DE-AC02-05CH11231. Curation tools are supported in part by the National Library of Medicine NLM R01LM013871.

